# Antitumor immunity in dMMR colorectal cancers requires interferon-induced CCL5 and CXCL10

**DOI:** 10.1101/2020.09.15.291765

**Authors:** Courtney Mowat, Shayla R. Mosley, Afshin Namdar, Daniel Schiller, Kristi Baker

**Affiliations:** Department of Oncology, University of Alberta, Edmonton, Canada; Department of Surgery, Royal Alexandra Hospital, Edmonton, Canada; Department of Medical Microbiology and Immunology, University of Alberta, Edmonton, Canada

**Keywords:** Colorectal cancer, tumor infiltrating lymphocytes, mismatch repair, chemokines, interferon (IFN)

## Abstract

Colorectal cancers (CRCs) deficient in DNA mismatch repair (dMMR) are heavily infiltrated by CD8+ tumor infiltrating lymphocytes (TILs) and are associated with a better prognosis than the majority of CRCs. The immunogenicity of dMMR CRCs is commonly attributed to abundant neoantigen generation due to their extreme genomic instability. However, lack of neoantigenic overlap between these and other CRCs necessitates study of antigen-independent mechanisms of immune activation by dMMR CRCs in order identify therapeutic strategies for treating MMR proficient CRCs. We show here using organoid cocultures and orthotopic models that a critical component of dMMR CRC’s immunogenicity is the activation and recruitment of systemic CD8+ T cells into the tumor epithelium by overexpression of the chemokines CCL5 and CXCL10. This is dependent on endogenous activation of the cGAS/STING and IFN signaling pathways by the damaged DNA in dMMR CRCs. These signaling pathways remain sensitive to exogenous stimulation in other CRCs, identifying an attractive therapeutic avenue for increasing TIL infiltration into normally immune resistant CRC subtypes. We have thus identified a key neoantigen-independent mechanism that underlies the ability for dMMR CRCs to recruit TILs into the tumor epithelium. Given that TIL recruitment is a prerequisite for effective tumor killing either by the endogenous immune system or in the context of immunotherapies, treatments that activate IFN-induced chemokine-production by tumor cells promise to improve the prognosis of patients with many different CRC subsets.

**Statement of Significance:** A critical component of antitumor immunity in dMMR CRCs is their ability to recruit T cells into the tumor epithelium as a prerequisite to tumor cell killing. This occurs because their extensive genomic instability leads to endogenous activation of cGAS/STING and overexpression of CCL5 and CXCL10.

## Introduction

Colorectal cancer (CRC) is a prevalent and often fatal disease initiated by either loss of the tumor suppressor APC or the DNA mismatch repair gene MLH1.^1,2^ Whereas APC inactivation promotes large scale DNA damage leading to chromosomal instability (CIN), MLH1 inactivation promotes hypermutability in the form of genome-wide point mutations due to deficient mismatch repair (dMMR). Although accounting for only 15-20% of CRCs, dMMR CRCs are of particular interest due to their potent immune-stimulating capacity and overall favorable patient outcome.^3^ This is commonly attributed to the high tumor mutation burden (TMB) induced by their hypermutability, which generates abundant neoantigens that can stimulate tumor infiltrating lymphocytes (TILs) to attack.^4^ Considerable effort is being invested in identifying immunogenic epitopes in the genomes of dMMR CRCs and there is no doubt that a high TMB contributes to immune activation by these CRCs.^5–7^ However, there is little overlap in the neoantigens of dMMR and CIN CRCs because of their different underlying mutational signatures.^8,9^ A narrow focus on neoantigen profiling of dMMR CRCs thus offers limited promise of discovering novel treatments to enhance immune activation in CIN CRCs. However, the lack of superior immunogencity or a better prognosis for dMMR tumors in other tissues indicates that a high TMB is neither necessary nor sufficient to explain these features in dMMR CRCs. There is thus a significant benefit to understanding the antigen-independent mechanisms of immune activation in dMMR CRCs that could be developed into therapies for a wider patient population.

Abundant TIL infiltration is an independent, positive predictive factor in many cancers including dMMR CRCs with immunologically “hot” tumors containing abundant infiltration of cytotoxic CD8+ T cells directly in the tumor epithelium.^10^ While the clinical significance of this pattern of TIL distribution within a tumor is clear, the processes regulating it are poorly understood. Critical outstanding questions relate to the source of these TILs, their activation or exhaustion level, their specificity for tumor neoantigens and the method of their recruitment. There is increasing evidence that tissue resident T cells, particularly CD8+ resident memory T cells (Trm), are critical mediators of antitumor immunity.^11,12^ In addition to the their localization within the diseased tissue and their inherently pre-primed state, Trm cells often express the appropriate surface receptors, such as CD103, for interacting with local epithelial cells. TILs in dMMR CRCs could thus arise from either an expansion of the resident intestinal intraepithelial lymphocyte population or recruitment of systemic T cells into the tumor. This is an important unresolved question since the former are inherently tolerized by their exposure to the suppressive mucosal immune environment and have a much higher activation threshold than CD8+ T cells recruited from systemic circulation.^13,14^ Furthermore, the mechanisms for recruiting these two immune populations will differ substantially.

Immune cell recruitment is commonly regulated through a complex network of chemokines that guide cell trafficking throughout the body.^15,16^ T cells in particular are responsive to a set of chemokines whose production is controlled by interferon (IFN) signaling.^17,18^ In most cases, this is initiated by intracellular viral infection and detection of pathogenic nucleic acids by intracellular DNA and RNA sensing systems such as cGAS/STING and AIM2, and TLR3.^19^ Numerous reports indicate that dMMR CRCs overexpress IFN stimulated genes (ISGs) yet what role this plays in TIL infiltration remains unknown.^20,21^ Additionally, most of these reports have drawn their conclusions from gene expression analysis of whole tissue transcripts without correcting for the disproportionately higher amount of immune cells in dMMR CRCs.^22,23^ Since immune cells are abundant producers of IFNs and ISGs, it is unclear if the dMMR CRC cells themselves or the infiltrated immune cells are the primary source of high IFN signaling in these tumors.

In the current study, we used physiological isogenic model systems to study how loss of the DNA mismatch repair gene *MLH1* in CRC cells drives the antigen-independent activation and recruitment of CD8+ T cells. We show that this requires induction of a gene signature that drives overexpression of the interferon-dependent chemokines CCL5 and CXCL10. This in turn leads to preferential recruitment and activation of systemic, not local, CD8+ T cells into the tumor epithelium. While activation of this immunogenic gene signature is endogenous in dMMR CRCs, it can be exogenously induced in CIN CRCs via IFN treatment or the induction of DNA damage. Given that immune recruitment is necessary for subsequent antigen-specific T cell-mediated killing, our findings identify therapeutic targets that could induce T cell infiltration into CIN CRCs, thereby providing the best chance for TIL activation even by these neoantigen-low tumors.

## Materials and Methods

### Cell line generation and stimulation

MC38 murine CRC cells were purchased from Kerafast. Cells were grown in high glucose DMEM supplemented with 10% FBS, 1% penicillin-streptomycin, and 1% HEPES at 37°C with 5% CO_2_. Cells were transfected using Lipofectamine 2000 (Thermo) with the pSpCas9(BB)-2A-puro (px459) V2.0 plasmid (Addgene) containing CRISPR guide-RNAs (gRNAs) (Supplementary Table 1) targeting the *Mlh1* gene to model dMMR CRCs or the *Kras* gene to model CIN CRCs. Transfectants were selected with 2ug/ml puromycin and mutations were confirmed either by sequencing or Western blot. shRNA-mediated knockdown of CCL5 and CXCL10 in the dMMR MC38 cells was achieved using the pLKO.1 system (Addgene #10878) and containing the shRNA sequences in Supplementary Table 1.^24^ Stably knocked down cells were selected with 250 ug/ml hygromycin and knockdown was confirmed by Western blot.

For sequencing, genomic DNA was isolated from cell pellets using the Quick Genomic DNA Extraction kit (Truin Science). Primers possessing EcoRI and BamHI cut sites at their ends were used (Supplementary Table 1) with the Q5 High-Fidelity PCR Kit (Thermo) to amplify CRISPR target site regions. PCR products were purified using a GeneJet Gel Extraction Kit (Thermo Scientific) and then the ends were digested using EcoRI and BamHI (New England Biolabs). PCR products were subcloned into the pUC19 plasmid (Addgene) and sequenced.

### Cancer cell stimulations

MC38 CRC cells were seeded into plates 24h before treatments as indicated in the figure legends. The following treatments were used: 10uM fludarabine, 50uM STAT3 Inhibitor VI (S3I-201), 10uM carbonyl cyanide 3-chlorophenylhydrazone (CCCP), 9ug/ml 2’,3’-cGAMP, 1uM 5-fluorouracil (5FU), 10uM N-methyl-N’-nitro-N-nitrosoguanidine (MNNG) (all from Sigma) or phosphorothioate oligo (IDT) or 1000U/ml IFNb1 (Thermo). After the indicated incubation time, cells supernatant or cells were harvested as indicated below.

### Mouse CRC experiments

C57BL/6 wildtype mice originally purchased from Charles River were bred and maintained in the Cross Cancer Institute vivarium. Mixed groups of male and female mice between the age of 6-20 weeks old were used for experiments. All animal work was approved by the University of Alberta Animal Care and Use Committee.

Orthotopic CRC experiments were performed by injecting 1×10^5^ MC38 CRC cells in 50ul PBS into the wall of the descending colon using a flexible needle (Hamilton) inserted through the working channel of a Wolfe endoscope and visualized via the ColoView imaging system (Storz).^25^ Orthotopic tumour growth was monitored by endoscopy and tumours were harvested after 2-3 weeks. Resected tumours were minced and digested in enzyme cocktail (RPMI containing 0.5mg/ml collagenase IV, 10μg/ml DNaseI, 10% FBS, 1% penicillin-streptomycin and 1% HEPES buffer) for 30 minutes at 37°C in a shaking incubator.^26^ Fragments were rigorously pipetted to dissociate, filtered through a 100μm cell strainer, washed and stained for flow cytometry.

Subcutaneous CRC experiments were performed by injecting 5×10^5^ MC38 CRC cells in 100ul PBS into the hind flank. Tumors were harvested after 2-3 weeks and digested as above. To separate the cancer and immune cells in the tumors, dissociated cells were resuspended in 40% percoll (GE Healthcare), overlaid onto 80% percoll and centrifuged at 500g (with brake off) for 30 minutes at room temperature. The top layer (tumour cells) and the interface (immune cells) were collected into separate tubes, washed, and processed for RNA isolation as below.

### Mouse and human CRC-derived organoids

Murine organoids from either the normal colonic epithelium or colorectal tumors induced by repeated doses of azoxymethane (10 weekly doses of 10mg/kg azoxymethane) were generated as described previously and cultured as below.^27,28^

Resected human CRC tumors were collected in HBSS within 10 min of devitalization. Tumors were processed as described previously.^28^ In brief, tumors were dissociated for 1h in DMEM with 2.5% FBS, 75U/ml collagenase XI, 125ug/ml dispase II (Sigma). Following filtration, cells were plated at 500-1000 per well in growth factor reduced Matrigel (Corning) and cultured in basal crypt media (Advanced DMEM/F12containing 10% FBS, 2mM glutamine, 10mM HEPES, 1mM N-acetylcystein, 1X N2 supplement, 1X B27 supplement, 10mM nicotinamide, 500nM A83-01, 10uM SB202190, 50ng/ml EGF) (ThermoFisher) mixed 1:1 with conditioned supernatant from L-cells expressing Wnt3a, R-spondin and noggin (ATCC #CRL-3276).^29^ All work with human samples was approved by the Health Research Ethics Board of Alberta.

Primary dMMR mouse or human organoids were generated using lentiviral transduction as described previously.^25,30^ Lentivirus was prepared as previously described using the pLKO.1 system (Addgene #10878) and containing the shRNA sequences in Supplementary Table 1.^24^ Lentivirus was concentrated 100X by ultracentrifugation. Organoids were pretreated for 4-5 days with 10mM nicotinamide to enrich for stem cells. Organoids were dislodged from the plate by pipetting and then treated for 5min at 37°C with TrypLE Express (Life Technologies). Organoids were mixed with lentivirus along with 8ug/ml polybrene and 10uM Y27632 (Sigma) and seeded into a 96-well plate. The plate was centrifuged for 60 min at 600g at 32°C and then incubated at 37°C for 6 h. The organoids were then embedded in Matrigel and cultured in media containing 100ug/ml hygromycin to select for successful transduction. Gene knockdown was verified by Western blot.

For stimulations, equal numbers of organoids were plated in Matrigel, cultured for 3 days and then treated as indicated in the figure legends. To harvest, organoids were resuspended in ice cold Cell Recovery Solution (CRS) (Corning) and incubated for 10min on ice to dissolve Matrigel. CRS was diluted 4-fold and then spun down. Pellets were processed for RNA or protein isolation.

### Flow cytometry

Flow cytometry staining was performed using the antibodies in Supplementary Table 2 in addition to the Zombie Aqua viability stain (BioLegend) and the Foxp3 Transcription Factor Staining Buffer Set (eBioscience). Flow cytometry was performed on a CytoFlex S cytometer (Beckman Coulter) with subsequent analysis using FlowJo (BD Biosciences).

### T cell activation and migration assays

CD8+ T cells were isolated from the spleens or mesenteric lymph nodes (MLNs) using the EasySep Mouse CD8+ T Cell Isolation Kit (StemCell Technologies). Where indicated, T cells were first expanded by incubation at a 5:1 ratio with bone marrow derived dendritic cells (BMDCs) that had been pulsed with tumor lysates (100ug/ml with 1×10^6^ BMDCs) for 30min and then irradiated (20 Gy). For direct coculture experiments, tumor cells were first incubated with the indicated treatment and this was removed before adding CD8+ T cells at a 5:1 T cell:tumor cell ratio for the indicated time. For chemokine blocking experiments, the antibodies (Supplementary Table 2) were present throughout the coculture at 2ug/ml. For migration assays, conditioned supernatants (DMEM + 10% FBS) were collected from CRC cells treated as indicated in figure legends. Tissue culture inserts with 5.0um pores (Sarstedt) were precoated with 0.8mg/ml Matrigel for 2h at 37°C and rinsed 2X with warm media. 1×10^5^ CFSE-labeled CD8+ T cells were added to the inner portion of the insert in 100ul media containing 2% FBS. 500ul of conditioned supernatants were added to the bottom well of the insert. For chemokine blocking experiments, antibodies were added to the media in the lower chamber 30min prior to T cell addition and were present during the entire assay. T cells were allowed to migrate for 3h and then cells in the inner insert and bottom well were collected and quantified by flow cytometry. % migrated cells were calculated as follows: % migrated cells = (# migrated cells) / (# input cells + # migrated cells).

### Western blotting

Protein was isolated in lysis buffer (50mM Tris-HCl, 150mM NaCl, 50mM sodium pyrophosphate, 1mM EDTA, 05% NP40, 1% Triton X-100) containing 1mM sodium orthovanadate, and 1x protease inhibitor (Sigma-Aldrich). Protein was quantified using the Pierce™ BCA protein assay kit (Thermo). 5 or 10ug of protein was loaded per lane and of SDS-PAGE gels and transferred to nitrocellulose membranes. The antibodies used are listed in Supplementary Table 2. Bands were visualized using the ECL Prime Western Blotting Detection Reagent (GE Healthcare Amersham).

### RNA and qPCR

RNA was extracted using Trizol and reverse transcribed using the High-Capacity cDNA Reverse Transcription Kit (Thermo). qPCR reactions were set up using the primers indicated in Supplementary Table 1 and POWRUP SYBR Master Mix (ThermoFisher). qPCR was performed on the QuantStudio6 real-time PCR system (Applied Biosystems).^118,119^

### Microarray

Microarray was performed on in vitro cultures of dMMR and CIN MC38 CRC cells using the SurePrint G3 Mouse Gene Expression v2 Microarray (8×60K) (Agilent) system. Biological replicates collected independently of each other were run in parallel for each sample. Gene Ontology Enrichment analysis was performed using PANTHER (http://pantherdb.org).^31^ Interferon Stimulated Genes (ISGs) were defined as previously described.^32^

### scRNAseq

Live cells were isolated from orthotopically grown dMMR and CIN MC38 CRC tumors. Each sample represents pooled cells from 5 mice per group. Viable CD45+ cells were enriched using magnetic selection (StemCell Technologies) and submitted to the University of Alberta High Content Analysis Core for processing with a Single Cell Immune Profiling kit (10X Genomics). The data was processed using the Seurat package for R (v3.0) as described previously.^33^ Briefly, cells with either a unique gene count over 2,500 or less than 200 as well as genes found in fewer than 3 cells were filtered out. Principle component analysis (PCA) was used to visualize and explore these datasets and we used the default settings of the RunTSNE function to summarize the PCA with tSNE dimensionality. Cell clusters in the two-dimensional representations were annotated to known cell types using FindAllMarkers for all clusters. We generated expression matrices with the Cell Ranger pipeline (10X Genomics) and converted them to .cloupe files in order to create two-dimensional tSNE plots. Gene Ontology Enrichment analysis was performed using PANTHER (http://pantherdb.org).^31^ Interferon Stimulated Genes (ISGs) were defined as previously.^32^

### The Cancer Genome Atlas analysis

Human RNA sequencing data and DNA sequencing data (Illumina HiSeq RNASeqV2) from the Colorectal Adenocarcinoma dataset from The Cancer Genome Atlas (Nature 2012, and PanCancer Atlas) for CRC were downloaded from cBioPortal for Cancer Genomics (https://www.cbioportal.org/).^34,36^

### Statistical analysis

Prism (GraphPad Software Inc.) was used for statistical analysis. Gene expression analysis was processed by log2 transformation and resulting data evaluated for Gaussian distribution. Comparisons of two unpaired groups was made by two-tailed Student’s t-test for normal data, or Mann-Whitney for non-parametric tests. For three or more groups with two parameters, two-way ANOVA or multiple t-test procedures were used as appropriate, for data with Gaussian distribution. Analysis of responses to multiple stimuli of a single cell type or of cells from a single donor were analyzed using paired tests. Post-hoc analysis to correct for multiple comparisons and detect differences between groups was by the two-stage linear step-up procedure of Benjamin, Krieger and Yekutieli with false discovery rate <0.05. A two-sided probability (p) of alpha error less than 0.05 defined significance.

## Results

### Mismatch repair deficiency in CRC induces a chemokine signature associated with CD8+ T cell recruitment

Immune-associated genes are well known to be overexpressed in dMMR CRC but few studies have investigated the functional significance of these genes for the antitumor immune response.^1,3^ Furthermore, most reports are based on whole tissue transcript analysis, making it difficult to determine the relative contribution of the CRC cells themselves to the expression of immune genes in the tissue.^23^ In seeking to better understand the factors governing TIL recruitment into dMMR CRCs, we examined expression of various chemokines in data obtained from The Cancer Genome Atlas PanCancer Atlas data set.^35^ We noted that two chemokines in particular that are known to regulate CD8+ T cell trafficking, CCL5 and CXCL10, were more highly expressed in dMMR CRCs than their CIN counterparts (Fig.1a).^22,37^ To determine if the dMMR CRC cells themselves were the primary source of these chemokines, we generated dMMR and control CIN variants of the MC38 CRC cell line by mutating *Mlh1* or *Kras*, respectively (Supplementary Fig.1). This led to the selective upregulation of chemokines CCL5 and CXCL10 specifically in the dMMR CRC cells (Figs.1b,c). In order to verify that the dMMR and CIN MC38 cells truly modeled their human CRC counterparts, we subcutaneously injected these cells into the flanks of immunocompetent wild type C57BL/6 mice and examined the immune profile of the resulting tumors. Consistent with human CRCs, dMMR tumors contained greater numbers of CD8+ T cells than CIN tumors (Fig.1d). Furthermore, whole tissue transcript analysis confirmed that the dMMR MC38 CRC tumors expressed greater amounts of CCL5 and CXCL10 compared to the CIN MC38 CRC tumors (Fig.1e). In contrast, dMMR CRCs did not contain more CXCL16, which is not known to be associated with CD8+ T cell recruitment (Fig.1b,e). These data indicate that loss of DNA mismatch repair in CRC triggers expression of chemokines specifically associated with TIL recruitment and that this may represent an important component of their immunogenicity.

**Figure 1.**
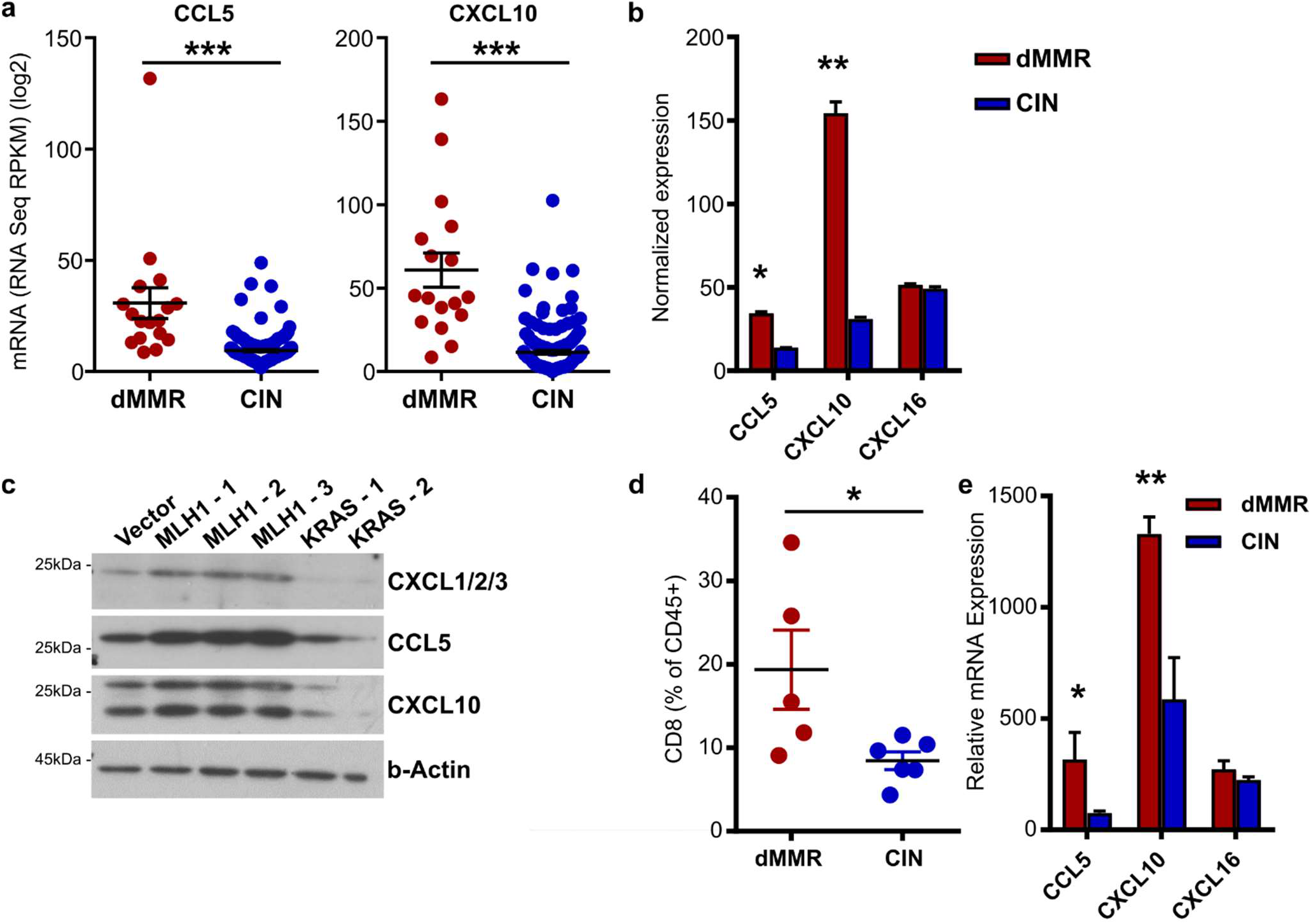
Mismatch repair deficiency in CRC induces a chemokine signature associated with CD8+ T cell recruitment. (a) Chemokine expression in dMMR CRCs and CIN CRCs from the TCGA PanCancer Atlas. (b) dMMR and CIN models of the MC38 mouse CRC cell line were created by mutating *Mlh1* and *Kras*, respectively. Chemokine expression in the cell lines was assessed by qPCR. n = 3 repeats. (c) Chemokine expression was analyzed by Western blotting lysates from 3 different clones of dMMR MC38 cells and 2 different clones of CIN MC38 CRC cells. Clones 1 were used in all subsequent experiments. n = 3 repeats. (d) Infiltration of CD8+CD3+ T cells in subcutaneously injected dMMR and CIN MC38 CRC tumors. n ≥ 4 mice per group, 4 repeats. (e) Chemokine expression was assessed by qPCR in whole tissue transcripts isolated from subcutaneously grown dMMR and CIN CRC. n ≥ 4 mice per group, 4 repeats. dMMR vs CIN: * p ≤ 0.05, ** p ≤ 0.01, *** p ≤ 0.005. See Supplementary Figure 1.

### Expression of a chemokine signature is essential for recruitment and activation of systemic CD8+ T cells by dMMR CRCs

Although the abundant infiltration of the dMMR CRC tumor epithelium by CD8+ TILs is a defining feature of these CRCs, the origin of these TILs or their mechanism of recruitment remains unknown.^2,38^ Of particular importance is determining if they originate from the systemic pool of CD8+ T cells or from the resident pool of immunologically tolerized T cells. In addition to their different activation states, these cells are expected to have fundamentally different recruitment mechanisms. We first determined if the CCL5 and CXCL10 produced by dMMR CRCs could recruit systemic CD8+ T cells using Transwell migration assays. Greater numbers of CD8+ T cells migrated towards supernatant conditioned by dMMR CRC cells than by CIN CRC cells and this was significantly inhibited with blocking antibodies against either CCL5 or CXCL10 (Fig.2a). We further confirmed the importance of CCL5 and CXCL10 in CD8+ T cell recruitment by knocking down CCL5 and CXCL10 in the dMMR MC38 cells. Supernatant from the chemokine deficient cells attracted significantly fewer CD8+ T cells than supernatant from the original dMMR CRC cells (Fig.2b). To explore the origin of TILs in dMMR CRCs, we next investigated how effectively our MC38 CRC variants could induce migration of CD8+ T cells from different sources. Whereas splenic CD8+ T cells were more attracted by conditioned media from dMMR CRC cells than media conditioned by CIN CRC cells, CD8+ T cells derived from the mesenteric lymph nodes (MLNs), which are the source of intestinal CD8+ T cells, were equally attracted by both tumor types (Fig.2c). Since both our in vivo experiments and patient data indicated that the TILs in dMMR CRCs are more activated than those in CIN CRCs, we further investigated where there was a difference in the capacity of dMMR CRCs to activate CD8+ T cells of different origins (Fig.1e). After coculturing splenic or MLN CD8+ T cells with the CRC cells for 24h, only the splenic CD8+ T cells were preferentially activated by dMMR CRC cells (Fig.2d). Collectively, these data suggest that the abundant TILs in dMMR CRCs result from selective recruitment and activation of systemic CD8+ T cells in a CCL5- and CXCL10-dependent manner.

**Figure 2.**
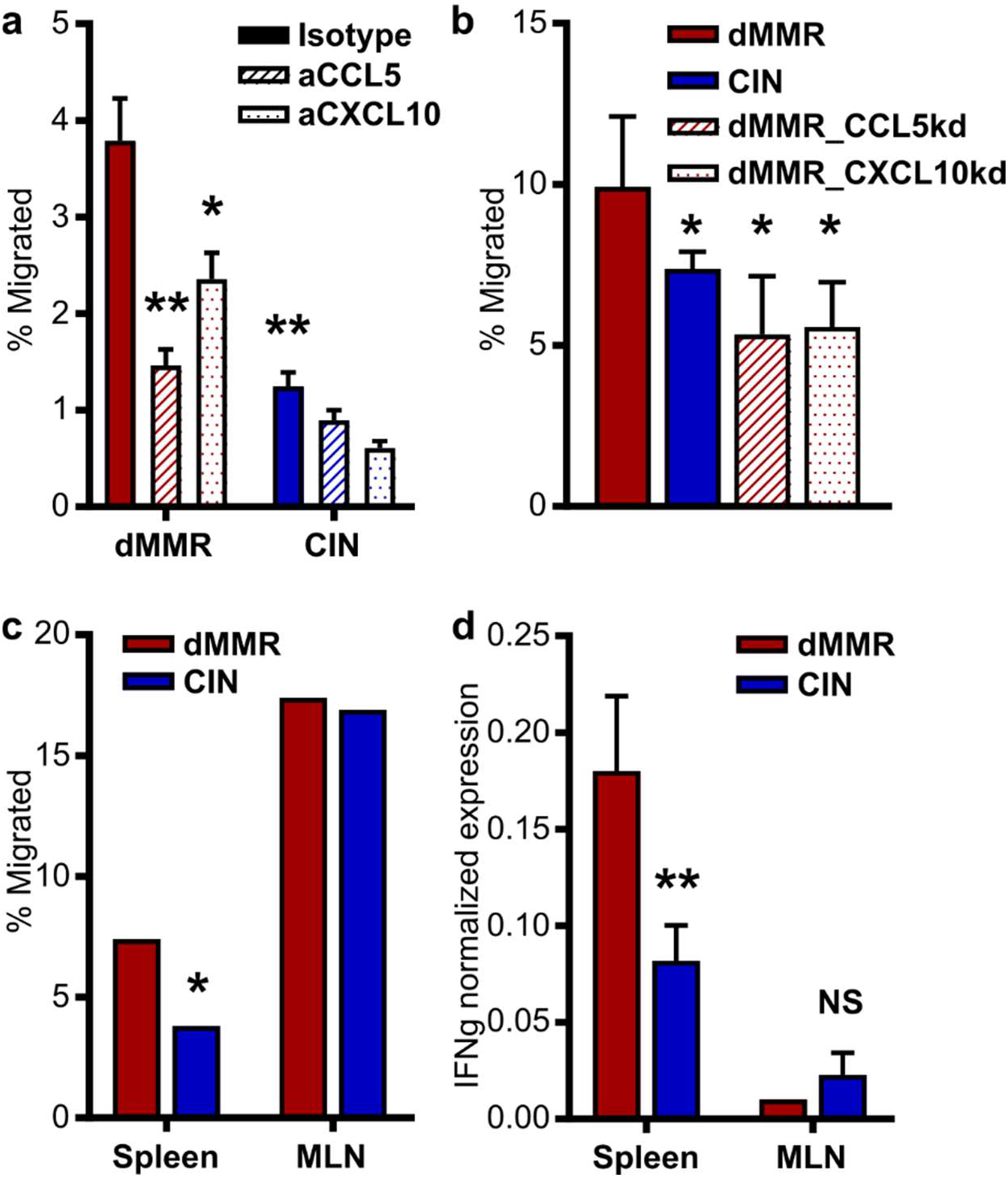
Expression of a chemokine signature is essential for recruitment and activation of systemic CD8+ T cell by dMMR CRCs. (a) CD8+ T cell migration through a Matrigel-coated 5.0 um Transwell insert towards supernatant conditioned for 24h by dMMR or CIN MC38 CRCs. Anti-CCL5, CXCL10 or isotype control antibodies were added at 2.0 ug/ml 30 min before T cells were added. Migration was carried out for 2 h and migrated cells were quantified by flow cytometry. n = 5 repeats. dMMR vs: * p ≤ 0.05, ** p ≤ 0.01. (b) CD8+ T cell migration towards supernatant conditioned by dMMR, CIN or dMMR cells deficient in either CCL5 or CXCL10. n = 3 repeats. dMMR vs: * p ≤ 0.05. (c) Migration of CD8+ T cells isolated from the spleen or mesenteric lymph nodes (MLN) towards conditioned supernatant from dMMR or CIN CRCs. Representative data from n = 4 repeats. dMMR vs CIN: * p ≤ 0.05. (d) Activation of CD8+ T cells isolated from the spleen or MLN and cocultured at a 5:1 ratio with dMMR or CIN CRC cells for 24 h. CD8+ cells were isolated and IFNg expression was quantified by qPCR. n = 3 repeats. dMMR vs CIN: ** p ≤ 0.01.

### Orthotopically implanted dMMR CRCs express more CCL5 and CXCL10 and recruit more CD8+ TILs than their CIN counterparts

T cell migration is a complex process involving tissue homing factors, adhesion to extracellular matrices of varied composition and cell-cell interactions. The only way to truly study such a complex process is using in vivo models with tumors growing in their endogenous environment. We thus orthotopically implanted the dMMR and CIN MC38 CRC cells directly into the colonic wall of wild type C57BL/6 mice in order to observe differences in immune cell infiltration in tumors growing in their native environment.^25,39^ dMMR CRC tumors consistently contained more CD8+ T cells than their CIN counterparts and these T cells expressed more IFNg, indicating greater activation (Fig.3a). Consistent with our findings relating to systemic CD8+ T cells as being the origin of TILs in dMMR CRCs, were failed to find differences in expression of the tissue residence marker CD103 on the TILs. Consistent with CRC patient data, transcript analysis of whole tumor tissue indicated that dMMR tumors expressed greater amounts of CCL5 and CXCL10, but not CXCL16, than did their CIN counterparts (Fig.3b). We did not observe differences in the infiltration by macrophages, dendritic cells, granulocytes, NK cells or CD4+ T cells indicating that recruitment by dMMR CRC was specific and was being directly orchestrated by the tumor cells themselves (Supplementary Fig.2). We furthermore confirmed the importance of CCL5 and CXCL10 at recruiting CD8+ TILs into dMMR CRCs by orthotopically injecting CCL5- and CXCL10-deficient CRCs. These chemokine-deficient tumors contained significantly fewer CD8+ T cells and far fewer of those that had infiltrated into the tumors expressed IFNg, indicating lower activation (Fig.3c).

**Figure 3.**
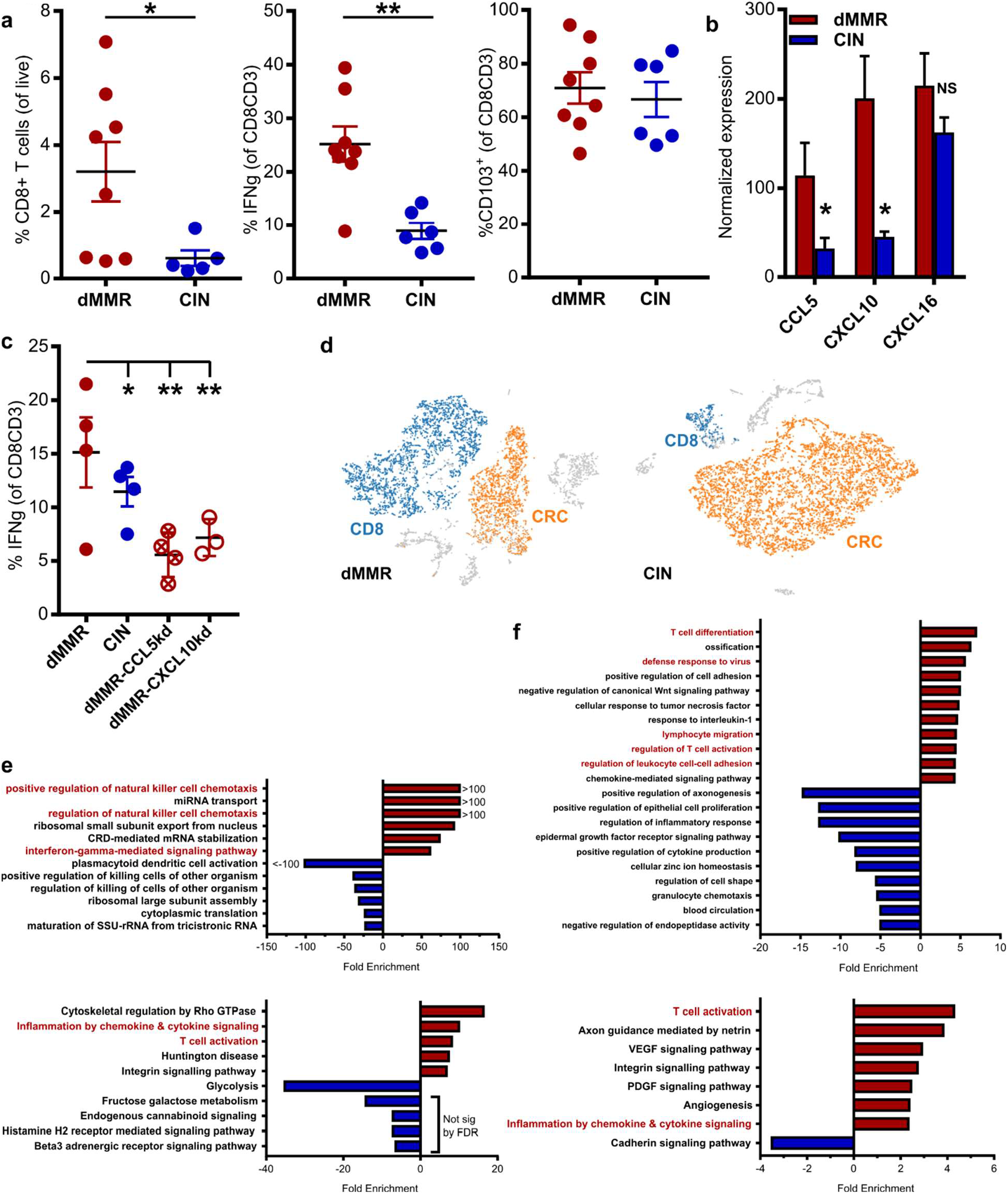
Orthotopically implanted dMMR CRCs express more CCL5 and CXCL10 and recruit more CD8+ TILs than their CIN counterparts. (a) Infiltration and activation of CD8+ T cells into dMMR and CIN CRCs grown orthotopically in the colon of wild type mice. 1.5×10^5^ CRC cells in 50 ul PBS were non-surgically injected into the colonic wall using an endoscope. Tumors were dissociated into single cell by enzymatic digestion and immune infiltration was assessed by flow cytometry. n ≥ 4 mice per group, 5 repeats. (b) Chemokine expression in whole tissue transcripts of orthotopically growth dMMR and CIN CRCs. (c) Activation of CD8+ T cells into orthotopically grown dMMR, CIN or dMMR cells deficient in either CCL5 or CXCL10. n = 4 mice per group, 2 repeats. (d-f) scRNAseq analysis of individual cells isolated from orthotopically grown dMMR and CIN CRCs. (d) tSNE plots were generated and differentially expressed genes were determined between dMMR and CIN CRCs in each of the indicated populations. (e-f) GO Enrichment Analysis of the most enriched biological processes (top) and signaling pathways (bottom) for CD8+ T cells (e) and MC38 CRC cells (f) identified in (d). n = 5 mice pooled per group. dMMR vs: * p ≤ 0.05, ** p ≤ 0.01. See Supplementary Figure 2.

To better understand the complex interplay between dMMR CRCs and their immune microenvironment, we performed single cell RNA sequencing (scRNAseq) on orthotopically grown dMMR and CIN CRCs (Fig.3d). CD8+ TILs within the dMMR CRCs were enriched in genes associated with T cell activation and chemokine-related signaling pathways in addition to several biological processes regulating chemotaxis (Fig.3e). Notably, Gene Ontology Enrichment Analysis revealed a significant enrichment of both T cell activation and chemokine related signaling pathways in the dMMR CRCs cells themselves in addition to a enrichment of several genes associated with the biological process of lymphocyte migration (Fig.3f).^31^ These data collectively support the hypothesis that chemokine secretion is a critical component of antitumor immunity dMMR CRCs and that the tumor cells themselves are central regulators of this process via production of CCL5 and CXCL10.

### Activation of endogenous IFN signaling in dMMR CRC leads to increased chemokine production

Chemokines CCL5 and CXCL10 are well known members of the Interferon Stimulated Genes (ISG) family, a group of genes induced by IFN and cGAS/STING signaling.^17,18^ scRNAseq analysis indicated significant differential expression of IFNA and IFNG-associated genes in dMMR CRCs, a finding we confirmed with qPCR of the CRC cells (Fig.4a,b). High ISG expression in dMMR MC38 CRC cells indicated that knocking out DNA mismatch repair increased either the endogenous activation of IFN signaling pathways within the cancer cells or the sensitivity of the cells to exogenous IFN. Given that we had not noticed consistently different expression of IFNB between our dMMR and CIN CRC cells and that there was not a significant source of exogenous IFN in the in vitro culture where differential CCL5 and CXCL10 gene expression were first noticed, we investigated the latter possibility first.

**Figure 4.**
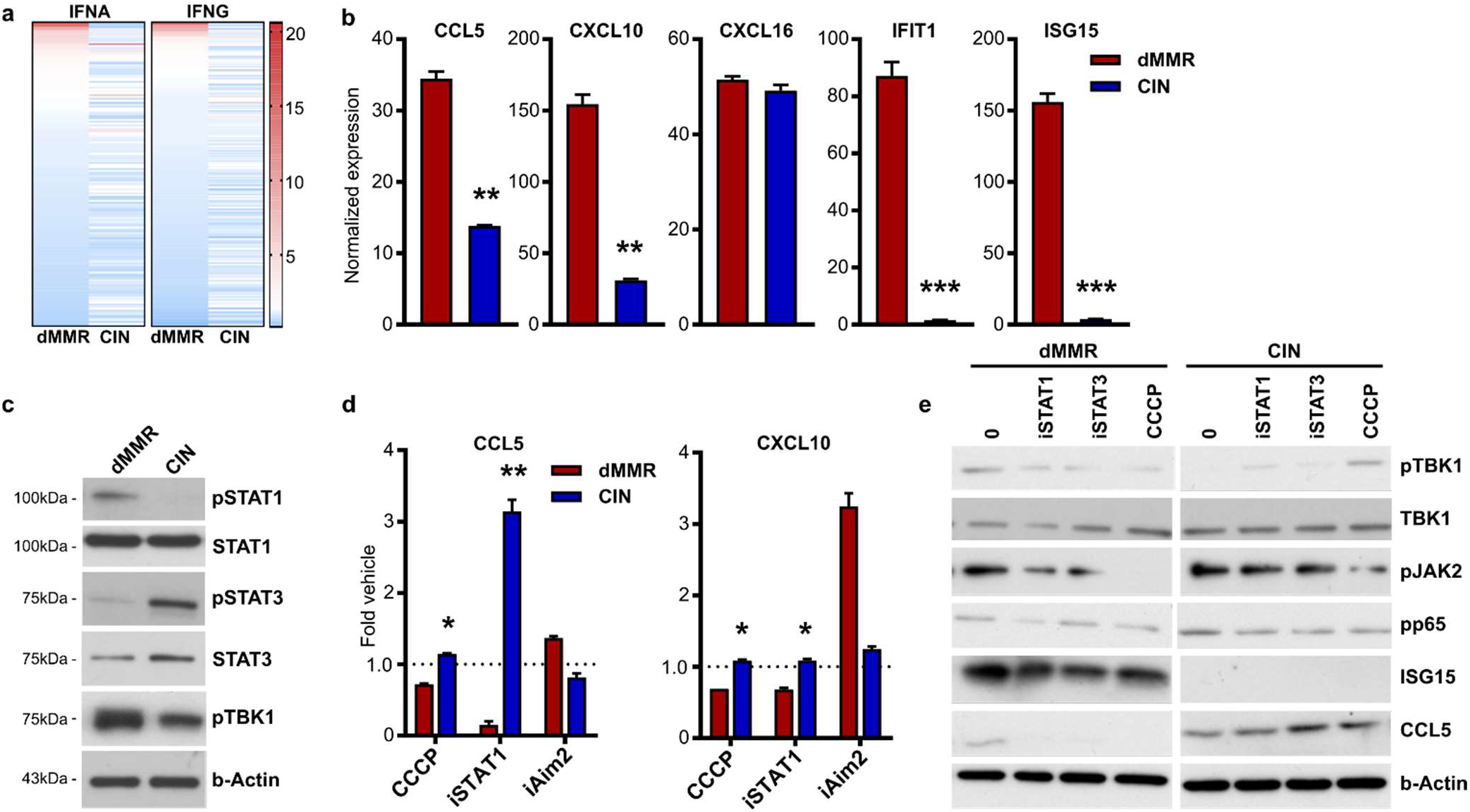
Activation of endogenous IFN signaling in dMMR CRC leads to increased chemokine production. (a) Expression of genes associated with IFNA and IFNG gene signatures in orthotopically grown CRC cells analyzed by scRNAseq (see Fig.3). (b) Baseline expression of key interferon stimulated genes (ISGs) or control genes in dMMR and CIN CRC MC38 cells. qPCR was performed on RNA isolated from unstimulated CRC cells grown in vitro. n = 4 repeats. (c) Baseline activation of proteins in the IFN and cGAS/STING signaling pathway in unstimulated whole cell lysates was determined in lysates from untreated dMMR and CIN MC38 CRC cells grown in vitro. n = 2 repeats. (d) Chemokine expression was determined by qPCR in dMMR and CIN CRCs treated for 24 h with inhibitors of STING (10 uM CCCP), STAT1 (10uM fludarabine) or AIM2 (9 ug/ml phosphorothioate oligo) or a vehicle control. n = 3 repeats. (e) Activation of IFN and cGAS/STING-associated signaling in cells treated as in (d) for 1 h. n = 2 repeats. dMMR vs CIN: * p ≤ 0.05, ** p ≤ 0.01, *** p ≤ 0.005.

We observed greater baseline activation of TBK1 and STAT1, but not STAT3, in dMMR CRC cells compared to CIN CRC cells (Fig.4c). TBK1 is a primary mediator of the cGAS/STING signaling pathway that drives innate interferon signaling and STAT1 is a downstream responder to activation of this pathway.^21^ Treating MC38 cells with inhibitors of STAT1 (fludarabine) or STING (CCCP) led to downregulation of CCL5 and CXCL10 selectively in dMMR but not CIN cells (Fig.4d). Since we did not observe such a decrease in chemokine production upon blocking the AIM2 nucleic acid sensing pathway, this does not seem to represent a generalized increase in cytoplasmic DNA sensing but rather specific activation of the cGAS/STING pathway. Consistent with this, both Fludarabine and CCCP also decrease endogenous activation of TBK1 and the STAT1-mediator JAK2 only in dMMR CRC cells (Fig.4e). These findings confirm that loss of MLH1 activates the endogenous IFN signaling pathway in dMMR CRC cancer cells, thereby increasing their ability to recruit CD8+ T cells into the tumor epithelium via the production of chemokines CCL5 and CXCL10.

### Both dMMR and CIN CRC cells remain sensitive to exogenous activation of IFN signaling despite differences in baseline IFN activation

IFNs are powerful inducers of immune-related signaling in both immune and non-immune cells such as epithelia.^18^ We next sought to determine if CIN CRCs have a fundamentally defective IFN signaling system that could explain our observations and contribute to their inability to recruit immune cells into the tumor microenvironment. Stimulation with the STING agonist cGAMP consistently upregulated CCL5, CXCL10 and ISG15, but not the non-ISG CXCL16, in both dMMR and CIN CRC cells, indicating that the STING signaling mechanism is intact even in the CIN MC38 cells (Fig.5a). Exogenous IFNB also consistently increased expression of ISGs in both CRC cell types, confirming that although quiescent at baseline, the mechanics of IFN signaling are intact in CIN CRCs (Fig.5b). This is particularly important given that immune cells infiltrating into a tumor can secrete large amounts of IFNs that could, in turn, activate antitumor immune gene programs in CRC cells. We furthermore observed that in the absence of exogenous IFN, blocking the IFNAR1 receptor substantially decreased ISG expression in dMMR CRC cells, pointing to an endogenous origin for the enhanced expression of ISGs in these cells (Fig.5c). We This led us to search for the mechanism behind such an increased sensitivity.

**Figure 5.**
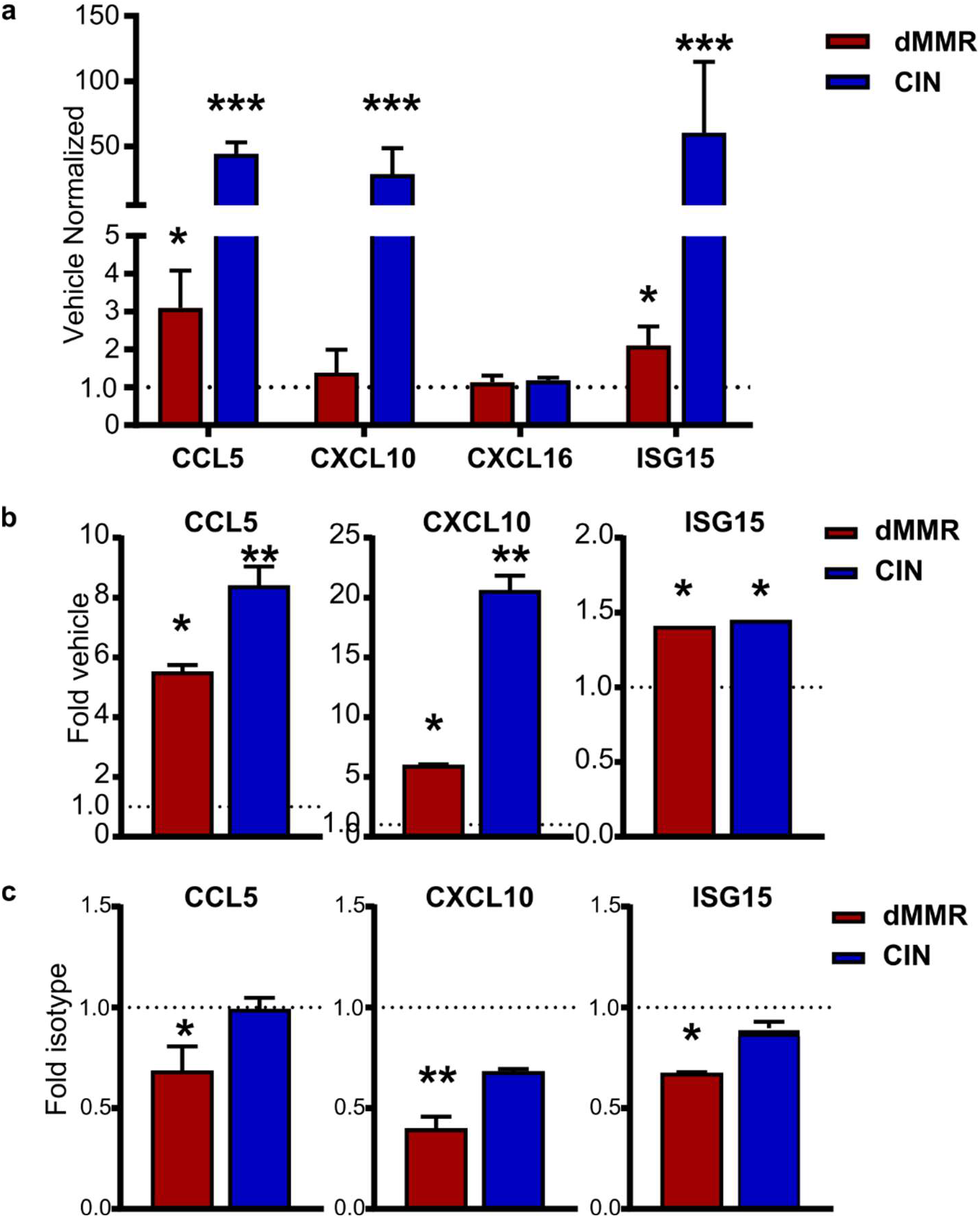
Both dMMR and CIN CRC cells remain sensitive to exogenous activation of IFN signaling despite differences in baseline IFN activation. (a) ISG upregulation induced by exogenous activation of cGAS/STING. Cells were treated with 9 ug/ml cGAMP for 24 h and change in gene expression was quantified by qPCR. n = 3 replicates. Vehicle vs: *** p ≤ 0.005. (b) ISG upregulation induced by stimulation with 1000 U/ml of IFNB for 24 h before RNA isolation and qPCR. n = 3 repeats. Vehicle vs: * p ≤ 0.05, ** p ≤ 0.01. (c) Endogenous activation of ISGs is decreased by an IFNAR1-blocking antibody. Cells were treated for 24 h with 10 ug/ml of anti-IFNAR1 or an isotype control before RNA isolation and qPCR. n = 2 repeats. Isotype vs: * p ≤ 0.05, ** p ≤ 0.01.

### Genetic instability induced by loss of DNA mismatch repair changes baseline and treatment-induced activation of endogenous IFN signaling in CRC cells

Loss of the DNA mismatch repair function in dMMR CRCs confers widespread genomic instability that renders the tumors hypermutable. A notable consequence of extensive genomic instability in cancer cells is the release of damaged DNA into the cytoplasm in the form of either free DNA or micronuclei.^21^ Since such cytoplasmic DNA can activate nucleic acid sensing pathways in the cells, a plausible explanation for high endogenous activation of cGAS/STING signaling in dMMR CRC cells is that these cells have alterations in endogenous cytoplasmic DNA due their extreme underlying genomic instability. Immunofluorescence staining with an anti-double stranded DNA (dsDNA) antibody confirmed that our cells did contain significant quantities of cytoplasmic DNA although we did not observe any difference in quantity between dMMR and CIN CRCs (Fig.6a). We thus reasoned that loss of mismatch repair could alter the composition of cytoplasmic DNA in a way that more strongly activated endogenous IFN signaling, leading to enhanced chemokine production. In order to investigate this, we made use of the compound N-methyl-N’-nitro-N-nitrosoguanidine (MNNG) to induce DNA damage that is corrected via the mismatch repair pathway.^40^ MNNG treatment decreased production of CCL5 and CXCL10 only in dMMR MC38 CRC cells that are not capable of repairing the induced damage (Fig.6b). This indicates that failure to repair DNA mismatches can directly alter chemokine secretion.

**Figure 6.**
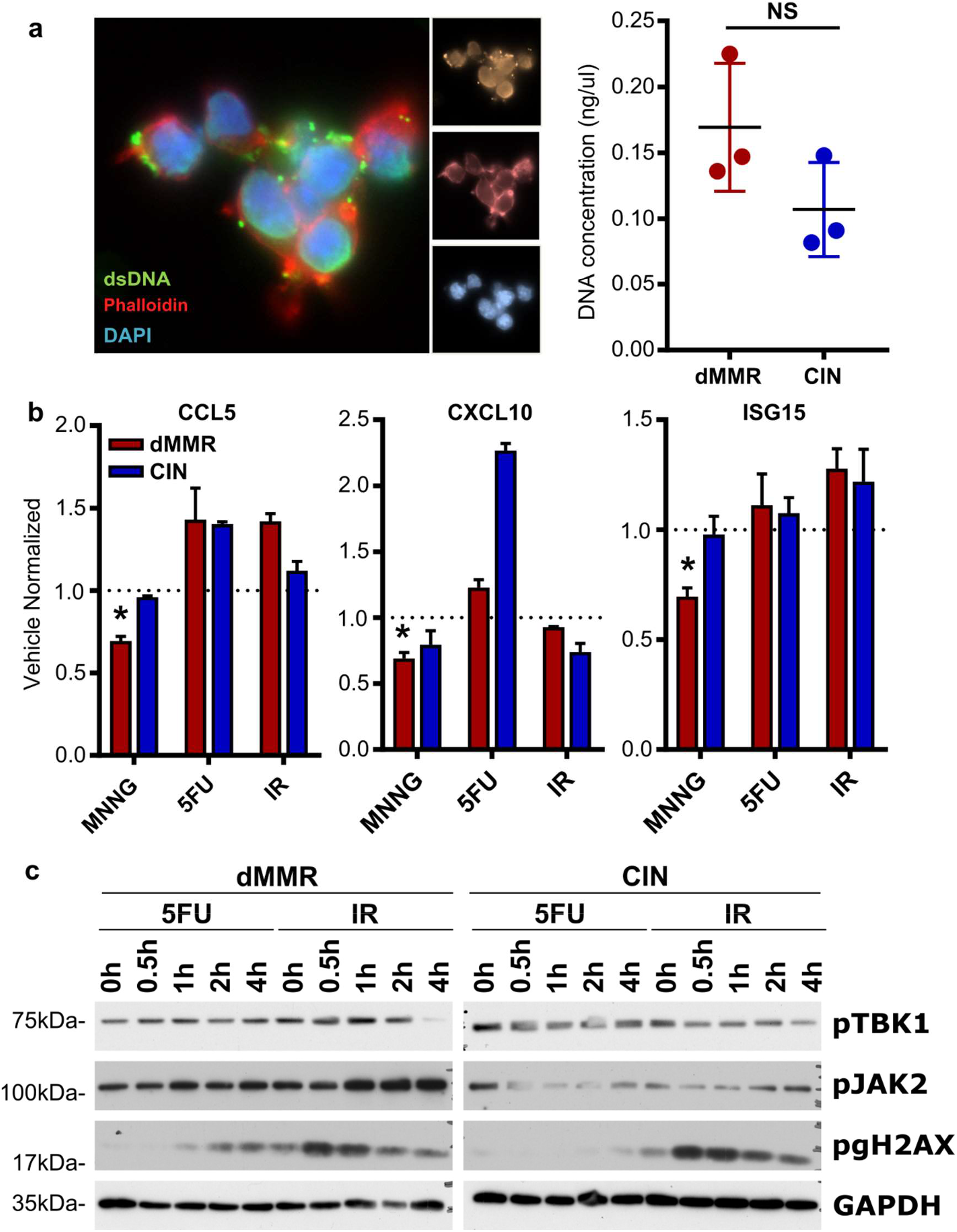
Genetic instability induced by loss of DNA repair changes baseline and treatment-induced activation of endogenous IFN signaling in CRC cells. (a) Cytoplasmic DNA was visualized by staining with an anti-dsDNA antibody (right). Cytoplasmic DNA was isolated from untreated dMMR and CIN MC38 cells and quantified by Qubit (left). n = 2 repeats. (b) dMMR and CIN MC38 CRC cells were treated with genotoxic agents (10uM MNNG, 1 uM 5FU, 10 Gy IR) for 24 h before RNA quantification and qPCR analysis. n = 4 repeats. (c) Cells were treated as in (b) for the indicated times before protein isolation and Western blotting. n = 3 repeats. dMMR vs CIN: * p ≤ 0.05.

Although MLH1 is the canonical mismatch repair protein, it also contributes to other DNA repair mechanisms such as base excision repair (BER) and homologous recombination (HR) ^41^ This is consistent with the longstanding observation that dMMR CRCs respond differently to many standard-of-care DNA damaging chemo- and radiotherapies than CIN CRCs.^42^ We tested whether increasing genomic instability induced by such treatments could alter chemokine production by exposing dMMR and CIN MC38 cells to either 5-fluorouracil (5FU) or ionizing radiation (IR). Both treatments equally upregulated CCL5 and CXCL10 in the dMMR and CIN CRC cells (Fig.6b). Treatment also increased the expression of other ISGs, and activation of TBK1 and JAK2, indicating generalized activation of cGAS/STING and IFN signaling (Fig.6c). These findings confirm that increased genomic damage generally increases chemokine expression via cGAS/STING. However, the findings also indicate that cGAS/STING-activating DNA associated specifically with defective DNA mismatch repair may have a differential capacity to endogenously upregulate signaling pathways leading to increased TIL recruitment and activation via CCL5 and CXCL10.

### Loss of DNA mismatch repair in CRC patient organoids upregulates chemokine production and IFN signaling

Cell line studies can be limited due to artifacts from long term cell culture. To determine if deletion of *MLH1* in primary CRC organoids also upregulated ISGs, we generated primary organoids from CRCs induced in wild type C57BL/6 mice through repeated doses of azoxymethane.^27^ Stably knocking down *Mlh1* in these organoids to model dMMR CRC indeed upregulated CCL5 and CXCL10, but not the non-IFN-induced chemokine CXCL16 (Fig.7a,b). Treatment of the dMMR organoids with the STING inhibitor CCCP decreased chemokine expression, confirming that chemokine induction in these primary cultures was also due to endogenous activation of the cGAS/STING signaling pathway (Fig.7b). Migration of CD8+ T cells towards supernatant conditioned by dMMR organoids was also suppressed to the same level as the control organoids by blocking antibodies against CCL5 and CXCL10, confirming the importance of these chemokines for T cell induced in the setting of primary dMMR CRC cells (Fig.7c).

**Figure 7.**
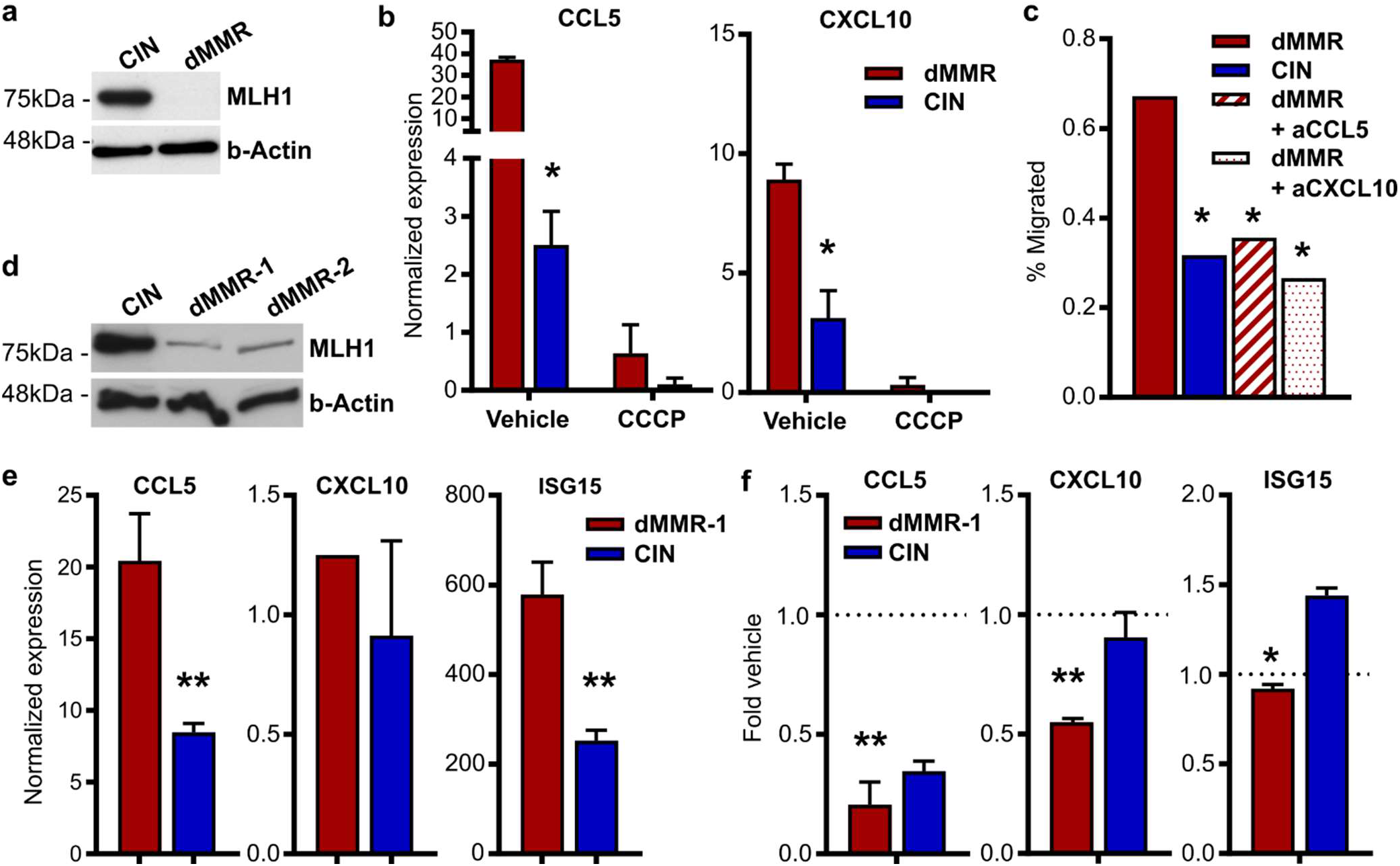
Loss of DNA mismatch repair in primary mouse and CRC patient organoids upregulates chemokine production and IFN signaling. (a) Baseline protein expression from primary mouse CRC organoids. CRC was induced by 10 weekly injections of 10 ug/kg azoxymethane followed by another 10 weeks without treatment. *Mlh1* was stably knocked down by shRNA to establish dMMR organoids by lentiviral transduction followed by hygromycin selection. Scrambled shRNA-transduced organoids were used as CIN controls. n = 2 repeats. (b) Reduced expression of ISGs following 24 h incubation of the dMMR and CIN organoids with 10 uM of the cGAS/STING inhibitor CCCP. n = 3 repeats. dMMR vs CIN: * p ≤ 0.05. (c) CD8+ T cell migration towards supernatant conditioned for 24 h by established dMMR or CIN CRCs. See Fig.3. dMMR vs: * p ≤ 0.05. (d-e) Baseline protein (d) and RNA (e) expression in organoids established from CRC patients. dMMR organoids were generated by stably knocking down MLH1 as in (b) with the appropriate shRNA and Scramble CIN control. dMMR vs CIN: ** p ≤ 0.01. (f) Inhibition of chemokine and ISG expression by treatment of human CRC organoids for 24 h with 10 uM CCCP. n = 2 patients, 3 repeats. Vehicle vs: * p ≤ 0.05, ** p ≤ 0.01.

To demonstrate the relevance of our findings to human CRCs, we generated organoids from resected CRC samples and stably knocked-down MLH1 in these organoids using lentiviral transduction (Fig.7d). This led to increased expression of the IFN stimulated genes CCL5, CXCL10 and ISG15, indicating that these chemokines are likely a significant contributor to TIL recruitment into dMMR tumors in CRC patients (Fig.7e). Treatment of the dMMR organoids with CCCP confirmed the importance of the cGAS/STING pathway to chemokine upregulation (Fig.7f). These data confirm that induction of CCL5 and CXCL10 via activated cGAS/STING signaling in dMMR CRCs is a direct result of loss of MLH1 and that activation of similar pathways in CIN CRCs represents a promising therapeutic approach for increasing TIL infiltration into this normally immune-resistant subset of CRCs.

## Discussion

Successful antitumor immunity requires the coordination of many processes, one of which is effective recruitment of TILs into the tumor epithelium. We show here that secreting TIL-recruiting chemokines is an essential immune-activating mechanism in dMMR CRCs. Specifically, genomic instability induced by defective DNA mismatch repair endogenously stimulates cGAS/STING signaling in dMMR CRCs. This activates the innate IFN signaling pathway thereby inducing expression of ISGs including the chemokines CCL5 and CXCL10. Blocking either cGAS or IFN sensing downregulates CCL5 and CXCL10 production by dMMR CRC cells while increasing genomic instability via DNA damaging agents increases their production. Similarly, neutralization of CCL5 and CXCL10 blocks migration of CD8+ T cells towards dMMR CRC cells, thereby precluding effective cytotoxic T cell killing of the tumors. Critically, cGAS/STING remains functional in CIN CRCs despite its low baseline activation, suggesting that therapeutically targeting this pathway could increase T cell trafficking into these normally immunologically silent tumors.

Abundant evidence links high TMB to tumor immunogenicity and activation of T cell-mediated killing. Paradoxically, some high TMB tumors fail to stimulate an endogenous antitumor immune response and remain refractory to checkpoint inhibition therapy.^8,43^ This includes tumors with defective MMR at other body sites as well as tumors rendered hypermutable by other mechanisms. One possible explanation for this is that mutational signatures caused by some processes are more immunogenic than others.^9^ Alternatively, hypermutation caused by different defects could selectively affect different gene families depending on their sequence or structure. This is consistent with the observation that, despite their mutator phenotype, specific genes are consistently found to be mutated in dMMR CRCs.^44^ The fact that this gene subset overlaps only partially with the subset of genes consistently mutated in dMMR endometrial cancers implies that site-specific selection factors may work in tandem with each mutational process to select for persistence of specific mutations from among a large body of randomly occurring passenger mutations. Our findings suggest that dMMR-induced hypermutability affects genes that regulate the production of chemokines such as CCL5 and CXCL10 and that this, in turn, generates strongly immunogenic tumors with a high TMB and enhanced ability to recruit cytotoxic T cells. Notably, our work implies that inducing the production of CCL5 and CXCL10 by tumor cells could recruit T cells into an environment that maximizes their likelihood of detecting even tumors with a low TMB such as CIN CRCs. Even low levels of tumor cell killing under these conditions could initiate development of effective antitumor immunity via processes such as epitope spreading.^45^ Our evidence that both dMMR and CIN CRCs remain sensitive to exogenous IFN further indicates that initial recruitment of CD8+ T cells, which are potent IFN producers, could establish a positive feedback loop where TILs induce tumor secretion of CCL5 and CXCL10, leading in turn to more TIL recruitment and activation.

Chemokines are potent but pleiotropic chemoattractants. Chemokine receptors often recognize multiple ligands and are broadly expressed on immune cells of different types.^15^ Our data indicate that both CCL5 and CXCL10 are selectively upregulated in dMMR CRCs. We consistently failed to find differences in expression of other chemokines such as CXCL16 that are not regulated by IFN. CXCL10 expression by cancers is consistently associated with effective antitumor immunity and CD8+ T cell regulation. In contrast, CCL5 expression has been linked to both pro- and antitumor immune responses. In particular, CCL5 can recruit tumor associated macrophages to the tumor microenvironment where they promote inflammation that facilitates cancer growth. ^37,46^ In addition, cancer cells themselves can upregulate CCR5, the CCL5 receptor, and respond to autocrine or paracrine section of CCL5. Indeed, blocking CCR5 with small molecule inhibitors such as maraviroc has shown some promise in clinical trials as an anti-cancer agent.^47^ Whether these compounds also inhibit antitumor immunity or TIL recruitment into tumors will be important to determine in future since this could compromise efficacy of the drug. In CRC, the majority of reports indicate that CCL5 expression is predictive of a positive patient outcome and productive antitumor immunity.^22,37^ Unfortunately, none of these reports have examined dMMR and CIN CRCs separately. Given that we observed a high correlation between CCL5 expression and mismatch repair deficiency that we observed in the data from TCGA as well as in our knockout cells, this could account for the apparent differential effect of CCL5 in CRC. Furthermore, consistent with the clinical literature on CRC in patients, we did not find any increase in tumor associated macrophages in orthotopically grown dMMR MC38 CRCs compared to CIN CRCs. Our blocking experiments indicate that CCL5 and CXCL10 additively or synergistically regulate CD8+ T cell trafficking and activation. This indicates that a gene signature with both of these chemokines could be predictive of patient prognosis and immunotherapy response even in CIN CRCs and particularly in the context of chemo- and radiotherapies that induce genomic instability.

The inherent ability of dMMR CRCs to stimulate antitumor immunity provides an important opportunity to learn the mechanisms necessary for this process. While there is an undeniable role for the high TMB content of these cancers, the lack of neoantigen overlap with CIN CRCs means that identifying other immune-stimulatory processes used by dMMR CRCs could provide more tractable therapeutic opportunities to treat other CRCs subsets. We have identified here a critical gene signature in dMMR CRCs consisting of endogenous IFN-induced production of CCL5 and CXCL10 and shown that this promotes antitumor immunity by recruiting systemic, non-tolerized, CD8+T cells directly into the tumor epithelium. This recruitment process appears to be independent of neoantigen production by the tumor cells, thereby establishing it as an achievable therapeutic target for low TMB CIN CRCs. Since TIL infiltration is a prerequisite for effective adaptive immune killing of tumor cells, our work provides not only a potential new screening tool for identifying patients likely to respond to immunotherapies but also establishes the basal criteria by which the immunogenic potential of other therapies, particularly DNA damaging agents, can be evaluated.

## Acknowledgements

The authors thank Rose-Marie Cornand, Dan McGinn, Cheryl Santos, Daming Li, Joaquín López-Orozco, Sharmin Sultana Sumi, Geraldine Barron, Dr. Xuejun Sun, Dr. Anne Galloway and Dr. Aja Rieger for technical support. Dr. Jatin Roper (Duke University) provided expert advice on orthotopic CRC protocols. This work was supported by funding from the Canadian Institutes of Health Research, the Natural Sciences and Engineering Research Council of Canada, the Canadian Foundation for Innovation, the Cancer Research Society and the University Hospital Foundation (KB).

