## Supplementary Tables and Figures for "Antitumor immunity in dMMR colorectal cancers requires interferon-induced CCL5 and CXCL10"

**Affiliations**

**Supplementary Table 1.**

| **Table 1. Primers** | | |
|---|---|---|
| **Primer Name** | **Primer Sequence (5'→3')** | **Purpose** |
| *Mlh1* A gRNA Forward | caccgGACGGTAGTGAACCGCATAGCGG | CRISPR |
| *Mlh1* A gRNA Reverse | aaacCCGCTATGCGGTTCACTACCGTCc | CRISPR |
| *Mlh1* B gRNA Forward | caccgGGTAGTGAACCGCATAGCGGCGG | CRISPR |
| *Mlh1* B gRNA Reverse | aaacCCGCCGCTATGCGGTTCACTACCc | CRISPR |
| *Kras* A gRNA Forward | caccgGCCGCCTGCCGAATCGAGCCCGG | CRISPR |
| *Kras* A gRNA Reverse | aaacCCGGGCTCGATTCGGCAGGCGGCc | CRISPR |
| *hMLH1*_053_shRNA Forward | AATTGTGTTCTTCTTTCTCTGTATTCTCGAGAATACAGAGAAAGAAGAACACTTTTTTTAT | shRNA |
| *hMLH1*_053_shRNA Reverse | AAAAAAAGTGTTCTTCTTTCTCTGTATTCTCGAGAATACAGAGAAAGAAGAACAC | shRNA |
| *Ccl5*_shRNA Forward | CCGGCGTGTTTGTCACTCGAAGGAACTCGAGTTCCTTCGAGTGACAAACACGTTTTTG | shRNA |
| *Ccl5*_shRNA_Reverse | AATTCAAAAACGTGTTTGTCACTCGAAGGAACTCGAGTTCCTTCGAGTGACAAACACG | shRNA |
| *Cxcl10*_shRNA Forward | CCGGTCCGGAATCTAAGACCATCAACTCGAGTTGATGGTCTTAGATTCCGGATTTTTG | shRNA |
| *Cxcl10*_shRNA_ Reverse | AATTCAAAAATCCGGAATCTAAGACCATCAACTCGAGTTGATGGTCTTAGATTCCGGA | shRNA |
| M13/pUC Forward | CCCAGTCACGACGTTGTAAAACG | Sequencing |
| M13/pUC Reverse | AGCGGATAACAATTTCACACAGG | Sequencing |
| *Mlh1* Seq Forward | GCGCGCGAATTCCCAAATCAAATGTCCGAGGGC | Sequencing |
| *Mlh1* Seq Reverse | GCGCGCGGATCCGTAGCAGGAGTTATTCGGCGT | Sequencing |
| *Ccl5* Forward | GCTGCTTTGCCTACCTCTCC | qPCR |
| *Ccl5* Reverse | TCGAGTGACAAACACGACTGC | qPCR |
| *Cxcl10* Forward | CCAAGTGCTGCCGTCATTTTC | qPCR |
| *Cxcl10* Reverse | GGCTCGCAGGGATGATTTCAA | qPCR |
| *Cxcl16* Forward | CTTCTGGCACCCAGATACCG | qPCR |
| *Cxcl16* Reverse | AGTTCCACACTCTTTGCGCT | qPCR |
| *Isg15* Forward | GGTGTCCGTGACTAACTCCAT | qPCR |
| *Isg15* Reverse | TGGAAAGGGTAAGACCGTCCT | qPCR |
| *Ifng* Forward | GGCAAAAGGATGGTGACATGA | qPCR |
| *Ifng* Reverse | ACCTGTGGGTTGTTGACCTC | qPCR |
| *Gapdh* Forward | CATGTTCCAGTATGACTCCA | qPCR |
| *Gapdh* Reverse | TGAAGACACCAGTAGACTCC | qPCR |

**Supplementary Table 2.**

| **Table 2. Antibodies** | | | |
|---|---|---|---|
| **Target** | **Fluorophore or Secondary** | **Purpose** | **Source** |
| b-Actin | Anti-rabbit IgG HRP | Western Blot | Cell Signaling (8457S) |
| MLH1 | Anti-rabbit IgG HRP | Western Blot | Abcam (ab92312) |
| Phospho-TBK1 (Ser172) | Anti-rabbit IgG HRP or Alexa 488 | Western Blot, Flow Cytometry | Cell Signaling (5483S) |
| TBK1 | Anti-rabbit IgG HRP | Western Blot | Cell Signaling (3504S) |
| CXCL1/2/3 | Anti-mouse IgG HRP | Western Blot | Santa Cruz (sc-365691) |
| CCL5 | Anti-rabbit IgG HRP | Western Blot | AbCam (ab189841) |
| CXCL10 | Anti-mouse IgG HRP | Western Blot | Santa Cruz (sc-374092) |
| Phospho-STAT1 (Tyr701) | Anti-rabbit IgG HRP | Western Blot | Cell Signaling (7649S) |
| STAT1 | Anti-rabbit IgG HRP | Western Blot | Santa Cruz (sc-464) |
| Phospho-STAT3 (Tyr705) | Anti-rabbit IgG HRP | Western Blot | Cell Signaling (9131S) |
| STAT3 | Anti-rabbit IgG HRP | Western Blot | Cell Signaling (12640S) |
| Phospho-NFκBp65 (Ser536) | Anti-rabbit IgG HRP | Western Blot | Cell Signaling (3033S) |
| Phospho-JAK2 (Tyr1007/1008) | Anti-rabbit IgG HRP | Western Blot | Cell Signaling (3776S) |
| ISG15 | Anti-rabbit IgG HRP | Western Blot | Cell Signaling (2743S) |
| Phospho-gH2AX | Anti-rabbit IgG HRP | Western Blot | Cell Signaling (9718S) |
| dsDNA | Anti-mouse IgG HRP | Immunofluorescence | Millipore (MAB1293MI) |
| CD3 | APCcy7 | Flow Cytometry | Biolegend (100222) |
| CD8 | APC | Flow Cytometry | Biolegend (100712) |
| CD4 | PerCPCy5.5 | Flow Cytometry | Biolegend (116012) |
| CD45 | PE | Flow Cytometry | Biolegend (103106) |
| CD69 | PeCy7 | Flow Cytometry | Biolegend (104512) |
| CD103 | FITC | Flow Cytometry | ThermoFisher (11-1031-85) |
| CD11c | PE | Flow Cytometry | Biolegend (117308) |
| CD11b | PECy7 | Flow Cytometry | Biolegend (101216) |
| H2kB | PE | Flow Cytometry | Biolegend (116507) |
| I-Ab | PerCPCy5.5 | Flow Cytometry | Biolegend (116416) |
| IFNg | PE | Flow Cytometry | Biolegend (505808) |
| Mouse IgG | HRP | Western Blot | Cell Signaling 7076S) |
| Rabbit IgG | HRP | Western Blot | Cell Signaling (7074S) |
| Rat IgG | HRP | Western Blot | Cell Signaling (7077S) |
| IFNAR1 | | Blocking | BioXcell (BE0241) |
| CCL5 | | Blocking | RnD Systems (MAB478-100) |
| CXCL10 | | Blocking | RnD Systems (MAB466-100) |
| Rat IgG Isotype | | Blocking | RnD Systems (MAB006) |

### Supplementary Figure 1

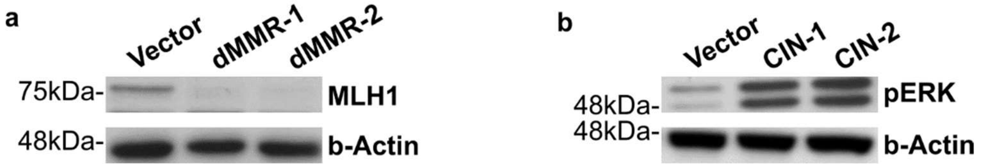


**Supplementary Figure 1. Confirmation of successful generation of dMMR and CIN MC38 CRC cell lines.** dMMR and CIN models of the MC38 mouse CRC cell line were created by mutating *Mlh1* and *Kras*, respectively. Protein expression was analyzed by Western blotting lysates from 2 different clones of dMMR MC38 cells and 2 different clones of CIN MC38 CRC cells. Clones 1 were used in all subsequent experiments. n = 3 repeats. See Figure 1.

**Supplementary Figure 2**

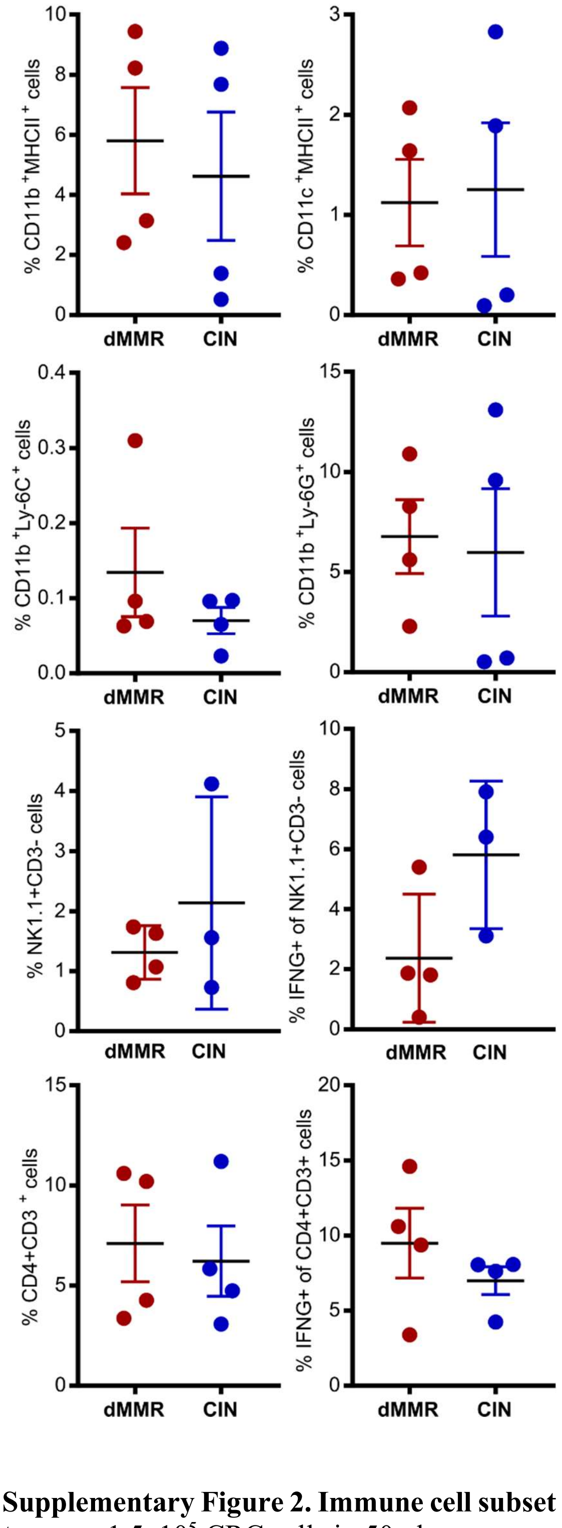


**Supplementary Figure 2. Immune cell subset frequencies in orthotopically implanted dMMR and CIN CRC tumors.** $1.5 \times 10^5$ CRC cells in 50 ul were non-surgically injected into the colonic wall using an endoscope. $n \geq 4$ mice per group, 5 repeats. Immune cell infiltrate was quantified by flow cytometry following tumor dissociation. See Figure 3.
